# On the imbalance between production and exploitation of marine fish assemblages: a case study from the Celtic Seas

**DOI:** 10.1101/2025.07.15.664907

**Authors:** Richard Law, Kennedy Edeye Osuka, Jon W. Pitchford, Michael J. Plank, James Rimmer, James Scott, Murray S. A. Thompson, Nicola D. Walker

**Affiliations:** University of York, York, UK; University of Liverpool, Liverpool, UK; University of Canterbury, Christchurch, New Zealand; Centre for Environment, Fisheries and Aquaculture Science, Lowestoft, UK

## Abstract

Sustainable management of marine ecosystems has to take into account the conservation of many species that are not themselves targets of fishing, in addition to those that are commercially exploited. Dynamic size-spectrum models suggest that fishing which leads species to have similar ratios of yield to production (i.e. similar exploitation ratios) is not sufficient to protect those that are rare: these species need to experience lower exploitation ratios. Here, the status of the demersal fish assemblage in the Celtic Seas is examined from the perspective of yield and production, using survey data collected over the period 2012 to 2016, and incorporating both common and rare species. The results give no evidence that rarer species have lower exploitation ratios than common ones. This suggests that current management methods are not not operating in a way that conserves the fish assemblage as a whole.

## 1 Introduction

The United Nations 2030 Agenda for Sustainable Development envisions a world in which humanity lives in harmony with nature, respecting biodiversity, and using natural resources sustainably in relation to their production (United Nations, 2023). Yet, in the case of oceans and seas, the goal of sustainable development (SDG 14) remains remote, and the signs are that humanity is mostly moving marine ecosystems in the wrong direction (Ye and Link, 2023). The goalposts of SDG 14.4 are themselves debatable, as they are aligned to Articles 61.3 and 119a of the Law of the Sea Convention (LOSC), which considers life in the oceans as a resource to be exploited on the basis of a maximum sustainable yield (MSY) (United Nations, 1982). This has the drawback that calculations of MSY are typically applied to a small number of commercially important species taken one at a time, and usually treated independently of the ecosystems in which they live, without taking into account the complex systems in which they are embedded. The MSY approach is questionable even for the commercially important species, because surplus production models have shown that, after aggregating to the ecosystem level, the MSY is less than the sum of the MSYs of the individual species (Link et al., 2012). Also, taking a precautionary approach, Spence et al. (2024) were unable to find precautionary reference points that in a multispecies setting would simultaneously maintain the biomass of nine species in the North Sea above levels at which biomass is considered threatened. Crucially neither the MSY nor the precautionary approach provides safeguards for rarer species important for conservation rather than commerce, yet still impacted by fishing activities.

Although fisheries management focuses primarily on the yields of commercially important species, there is a long history of interest in a more holistic approach to the management of marine ecosystems (Garcia et al., 2003; Jennings, 2005; Fogarty, 2014; Link, 2018). An important driver in this is the United Nations Convention on Biological Diversity (UNCBD) which, at an early stage, called for an ecosystem approach to conserve the structure and functioning of ecosystems, together with the biological diversity that underpins their functioning (5th Conference of the Parties (CoP), Decision V/6, Principle 5) (UNCBD, 2000). For the future of marine ecosystems, the challenge is to build a framework that meets the legal requirements of LOSC and the aspirations for conservation from UNCBD, such that exploitation and conservation can work together and prevent the loss of biodiversity (Garcia et al., 2016; UNCBD, 2023).

This paper builds a single framework within which the health of marine ecosystems can be assessed both from the perspective of expoitation and also from the perspective of conservation of biodiversity. At the heart of this is a recognition that marine ecosystems are complex systems, comprising intricate networks of organisms that live, grow and die through their interactions with one another. The basis for the framework is a relationship between the rate at which fish species generate biomass, i. e. the production rate (*P*), and the rate at which this production is removed as yield and bycatch (*Y*). There should be some relation between *Y* and *P* detectable across species living together in multispecies assemblages. *In extremis*, a species close to extinction has very low *P*, and will generate little yield before it disappears, whereas species with high *P* should be able to sustainably support some exploitation.

What should the relation between *Y* and *P* be in multispecies assemblages, and what actually happens in practice? If there is any yardstick, it is to work towards a constant ratio of *Y* to *P*, called the exploitation ratio *E*, often with the aim of achieving a value near to 0.5 (Patterson, 1992; Pikitch et al., 2012; ICES, 2024). For species of commercial importance (usually abundant species), the yields are subject to careful checks in the course of fisheries management. But less is known about the rate of removal of biomass from rarer species (i.e. of low abundance) in which fishing mortality is more a result of accidental bycatch. Moreover relatively little is known about the process of production itself, measured as the rate of accumulation of biomass through body growth of fish. The limited information available from theory warns that rare species may be more vulnerable and need lower exploitation ratios for sustainability than common species (Law and Plank, 2023). If rare species are driven towards extinction as a by-product of fishing activities, then exploitation fails the test of sustainability at the ecosystem level.

The theory relating *Y* and *P* in multispecies assemblages is technical (see Law and Plank, 2023). We therefore start with a brief explanation of the background: the theory suggests there are potential dangers of using a fixed exploitation ratio for ecosystem management, and an alternative approach is suggested. The remainder of the paper provides an example of the relationship between *Y* and *P* in an assemblage of demersal fish species in the Celtic Seas. There are two reasons for doing this. The first is to show that production rate can be estimated directly from survey data, and the paper proposes a novel method for doing so. The second is to see what can be learned about the health of the Celtic-Sea ecosystem from the *Y, P* relation. The results suggest that the *Y, P* relation is not consistent with conservation of rare demersal species. For conservation of the assemblage as a whole, the fishing mortality rates of species (rather than their yields) would need to be brought towards alignment with their production, a pattern of exploitation sometimes referred to as balanced harvesting (Garcia et al., 2016; Heath et al., 2017; Zhou et al., 2019; Law and Plank, 2023). This indicator of health could be a helpful harvest control rule at the ecosystem level (Garcia et al., 2016), meeting some requirements both of fisheries and of conservation.

## 2 Frameworks to connect production, yield and fishing mortality

Previous theoretical work (Law and Plank, 2023) used dynamic size-resolved ecosystem models to investigate sustainable ecosystem-level fishing strategies, and to explain how the concept of balanced harvesting needs careful interpretation, especially where this relates to the conservation of rare species. Real-world fisheries management, however, typically operates at the level of overall biomass quotas for individual species. Here we translate modelling insights into a framework which is consistent both with established management concepts and with available datasets. The key relationship is between the overall rate of production of biomass by species, and the rate at which this biomass is removed by fishing in a multispecies assemblage.

In shifting to familiar concepts of management, results from models structured by body size have to be condensed into unstructured species-level variables more commonly used in fisheries science. Thus the rate of production *P*_*s*_ is built from the rate at which fish of species *s* at a given body size accumulate biomass through body growth, by integrating over body size, and expressing the result as a mass per unit area per unit time (e.g. t km^*−*2^ yr^*−*1^) (Law and Plank, 2023, Eq. 4). We note that this aggregated measure of production is distinct from that used unstructured surplus-production models. *P*_*s*_ depends in part on the biomass density *B*_*s*_ of species *s*, obtained in the same way as *P*_*s*_, with dimensions mass per unit sea-surface area (e.g. t km^*−*2^) (Law and Plank, 2023, Eq. 3). Some part of the production is removed by fishing as yield *Y*_*s*_, and has the same dimensions as *P*_*s*_ (e.g. t km^*−*2^ yr^*−*1^) (Law and Plank, 2023, Eq. 5).

A simple way to think of the relationship between the aggregated variables *P*_*s*_, *Y*_*s*_, *B*_*s*_, is to assume that an assemblage is at dynamic equilibrium, such that the rate of loss of biomass is equal to the rate at which it is created in each species, through an assumed natural mortality rate *M*_*s*_ (yr^*−*1^) and fishing mortality rate *F*_*s*_ (yr^*−*1^):

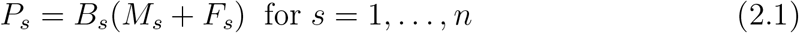

(Kolding et al., 2016), with *Y*_*s*_ defined as

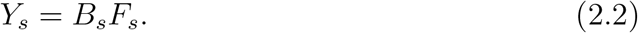

In reality (and in size-spectrum models), natural and fishing mortality are also functions of body size (Law and Plank, 2023), but this simplified version is sufficient to show some essential properties of the *Y, P* relationship. In particular, the ratio, *Y*_*s*_*/P*_*s*_, is the exploitation ratio *E*_*s*_, obtained from Eqs 2.1, 2.2:

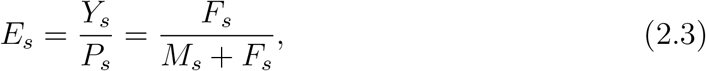

i.e. the proportion of production by species *s* removed as yield. It is a dimensionless measure, and is taken to indicate of overfishing if it rises above 0.4 to 0.5 (Patterson, 1992; Pikitch et al., 2012). It is used, for instance, in data-limited fish stocks in the guise of setting *F*_*s*_ = *M*_*s*_, equivalent to setting *E*_*s*_ = 0.5 (Lart, 2022; ICES, 2024) (see Discussion). Notice that, although this argument started by assuming an assemblage at equilibrium, the assumption does not apply to *E*_*s*_ itself, as this is simply a ratio of mortality rates. This means that it is entirely feasible for *E*_*s*_ of a species to remain unchanged, while, simultaneously, this species is collapsing to extinction (Law and Plank, 2023).

### 2.1 Fishing to a fixed exploitation ratio *E*

Working towards an exploitation ratio close to a fixed value *E*_*s*_ = *E* for every species would at first sight appear to be a rational solution to harvesting a multispecies assemblage (Kolding et al., 2016). From Eq. 2.3, it leads to a linear relationship between log*Y*_*s*_ and log*P*_*s*_:

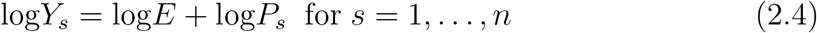

with a slope = 1 (Fig. 1a). This might be envisaged as a harvest control rule to apply to a whole assemblage of species, setting *F*_*s*_ proportional to the ratio *P*_*s*_*/B*_*s*_ for each species, where *E* is the constant of proportionality:

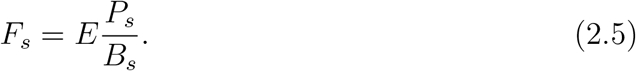

**Figure 1:**
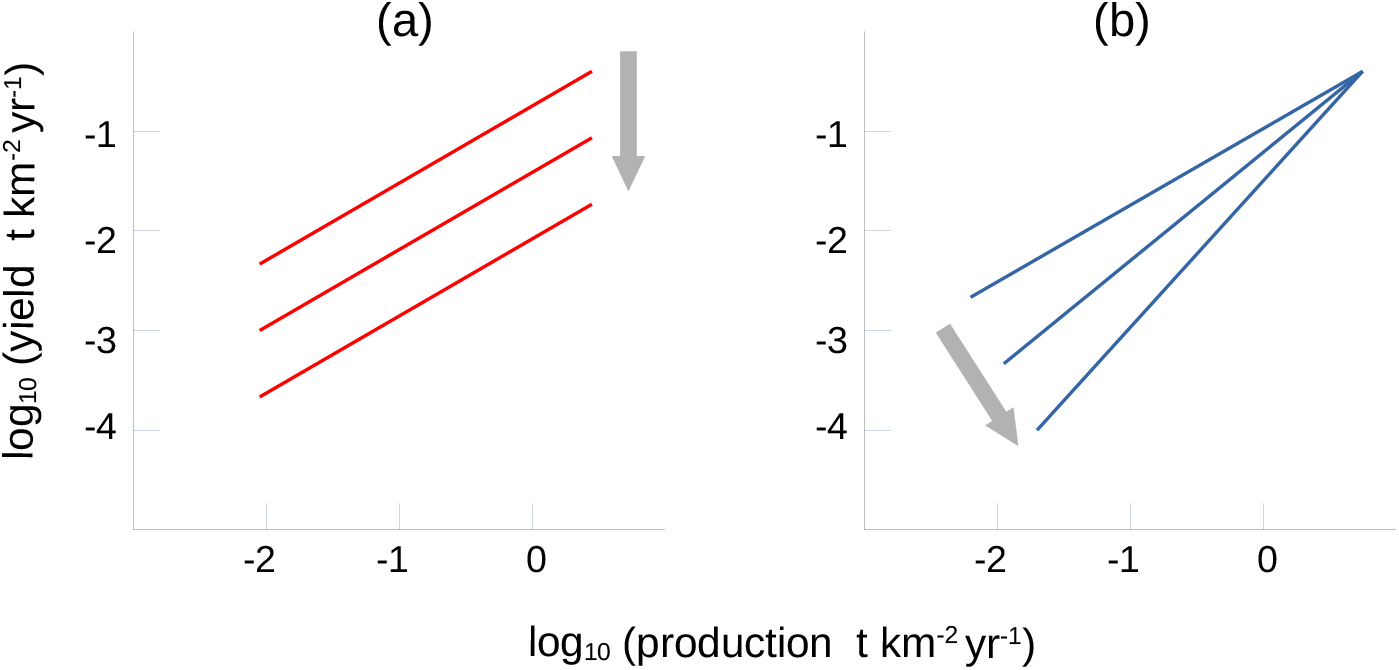
Relationships between yield *Y* and production *P*. (a) Fishing to a fixed exploitation ratio *E* brings the log*Y*, log*P* relation of a species assemblage near to a line of gradient 1, giving a family of parallel lines as *E* decreases shown by the arrow. (b) Reducing *E* for rare species increases the gradient, moving the assemblage towards balanced harvesting; this is needed to meet the dual objectives of generating yield from abundant species and conserving rare species.

However, setting *F*_*s*_s by the ratio *P*_*s*_*/B*_*s*_ also means there is no density dependence in fishing mortalities. This is evident from the fact that the dimensions of the ratio *P*_*s*_*/B*_*s*_ are 1/time (e.g. yr^*−*1^), and therefore independent of biomass density. In other words, rare species experience *F*_*s*_s as great as common ones. A likely outcome of the management strategy is that rare species needing more protection than common ones, are put on a path towards extinction. The destructive ecosystem-level consequences of such fishing were illustrated in Law and Plank (2023), where rarer species were shown to collapse even under a moderate fixed exploitation ratio. Put simply, while fishing to a constant exploitation ratio can deliver commercially viable yields of the more abundant fish species, theory suggests it is not compatible with the conservation of rare species that need more protection from fishing (Law and Plank, 2023).

### 2.2 Reducing the exploitation ratio *E* for rare species: balanced harvesting

The relationship between *Y* and *P* associated with a constant exploitation ratio *E* is just one of a family of possible harvest control rules for multipecies assemblages of the form

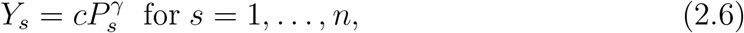

where the power parameter *γ* = 1 in the special case of a fixed exploitation ratio. Choosing a value of *γ >* 1 should reduce fishing pressure on species with low production rates relative to those with high production rates, giving a gradient *>* 1 in a log*Y*, log*P* plot (Fig. 1b).

This intuition is supported by modelling outputs from balanced harvesting (Law and Plank, 2023), where fishing mortality rates were set proportional to the production rates of each species, i.e. *F* ∝ *P*, rather than *F* ∝ *P/B* (Eq. 2.5). From Eq. 2.2, the yield under balanced harvesting becomes

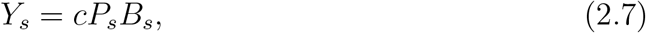

for some constant *c*. Eq. 2.7 is a special case of Eq. 2.6 because *P*_*s*_ and *B*_*s*_ are known to have a power relation in fish assemblages of the form *B* ∝ *P*^*α*^, with *α >* 0 (Heath et al., 2017). Anticipating our analysis of the Celtic Sea ecosystem in Section 3, Fig. 2 illustrates this relationship in its demersal fish species, the gradient in this instance being *α* = 0.865 (SE: 0.035) after log transformation. Thus Eq. 2.6 can be written as:

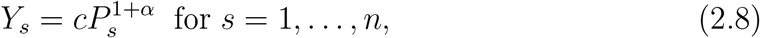

where the scaling constant (1 + *α*) is an emergent ecological property of an assemblage. This gives a linear relationship between log*Y*_*s*_ and log*P*_*s*_ similar to Eq: 2.4

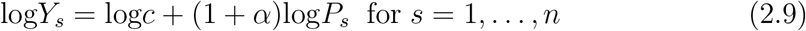

but with a different intercept and steeper slope (Fig. 1b).

**Figure 2:**
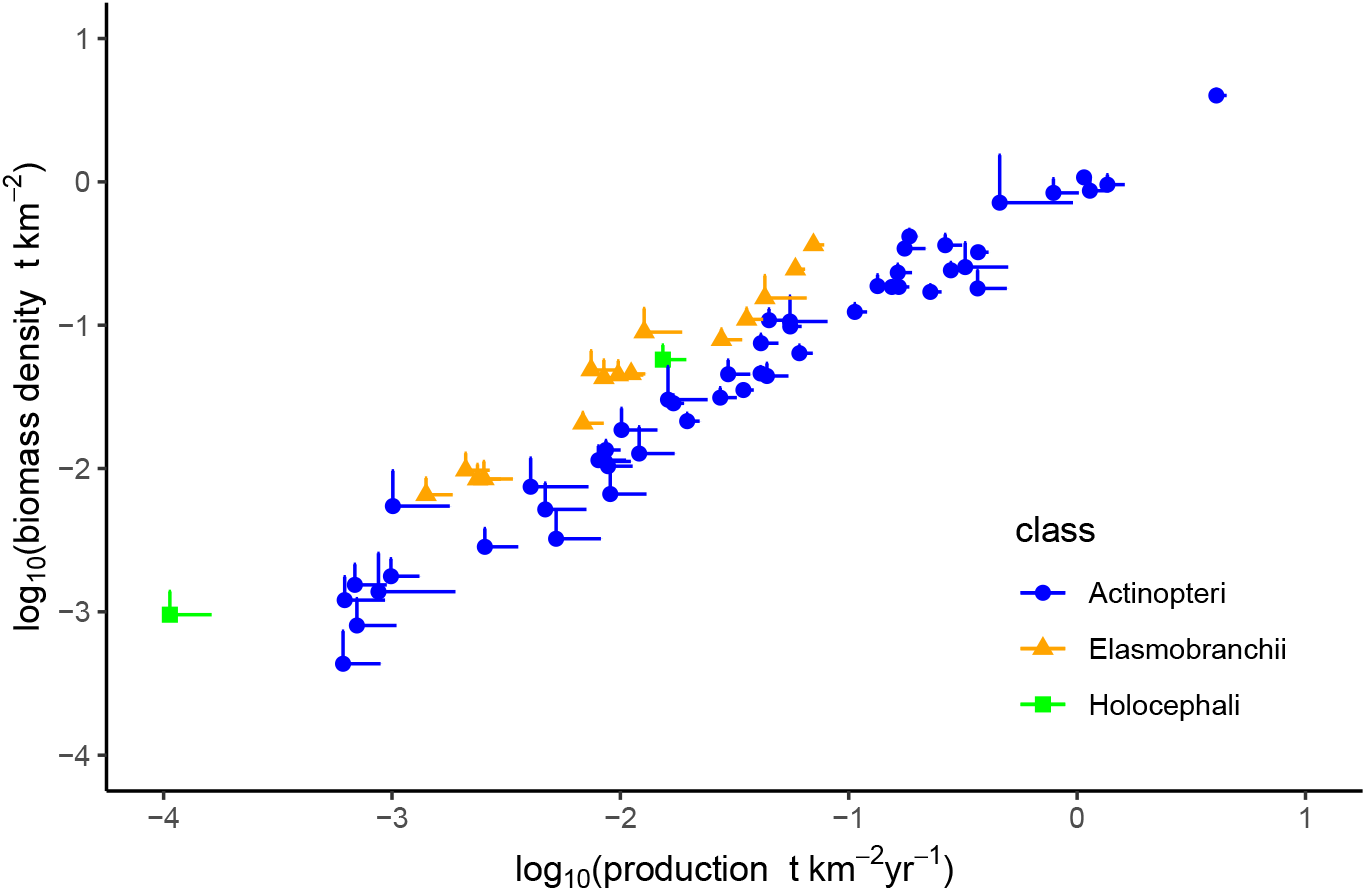
The relationship between production rate *P* and biomass density *B* of 67 demersal fish species in the Celtic Seas, estimated from otter- and beam-trawl survey data collected over the period 2012-16 from ICES Subarea 7b,c,e-k. Points are estimates for species with the upper 95 % confidence limits shown as bars. See Section 3 for a description of this ecosystem, and Appendix D, E and F for calculation of *B, P* and confidence limits, respectively.

In effect, moving fisheries in the direction of balanced harvesting, introduces density dependence into fishing mortality rates. This can be thought of as an ecosystem harvest control rule (ECHR) (see Section 4: Discussion). It allows commercial profitability of fisheries to be maintained by generating yield from common species with high production rates, and at the same time supports the conservation of rare species with low production rates.

## 3 A case study: Celtic Seas demersal fish assemblage

The arguments above from modelling dynamic fish assemblages suggest that the relationship between *Y* and *P* has some potential as an indicator of sustainability of exploitation of marine ecosystems. The log*Y*, log*P* plot can encompass the rarer species of concern for conservation in the same framework as common species of commercial importance. This comes with the caveat that such an assemblage is still no more than a subset of the overall biodiversity of the ecosystem.

To see what can be learned from the relationship between *Y* and *P*, we carried out a case study on an assemblage of demersal fish species in the Celtic Seas (ICES Subarea 7, divisions b, c, e-k) over the period 2012 to 2016 (Fig. 3). This ecosystem supports a major fishing industry (ICES, 2022). To obtain as much of the demersal fish assemblage as possible, the study was based on data from the International Bottom Trawl Survey (IBTS) which uses a standard otter trawl, and the Beam Trawl Survey (BTS). These surveys record all fish caught, which ensures that measures for all species are built on a common platform. Using this platform, we estimated measures of abundance, fishing mortality, yield and production rate of the species. We did this as a function of fish body length rather than fish age because (i) fish in the survey data were characterised by body length, and (ii) fishing mortality depends more on body size rather than on age. The surveys covered about 50 % of the area in ICES divisions 7b,c,e-k, but fishing effort data published by the European Union Scientific, Technical and Economic Committee for Fisheries (EU STECF) show that almost all of the commercial fishing effort (of the gears used here) was concentrated in the surveyed rectangles during the period 2012 to 2016. So there was a close spatial match between the surveys and the commercially fished areas. (See Appendix A for further details of the surveys.)

**Figure 3:**
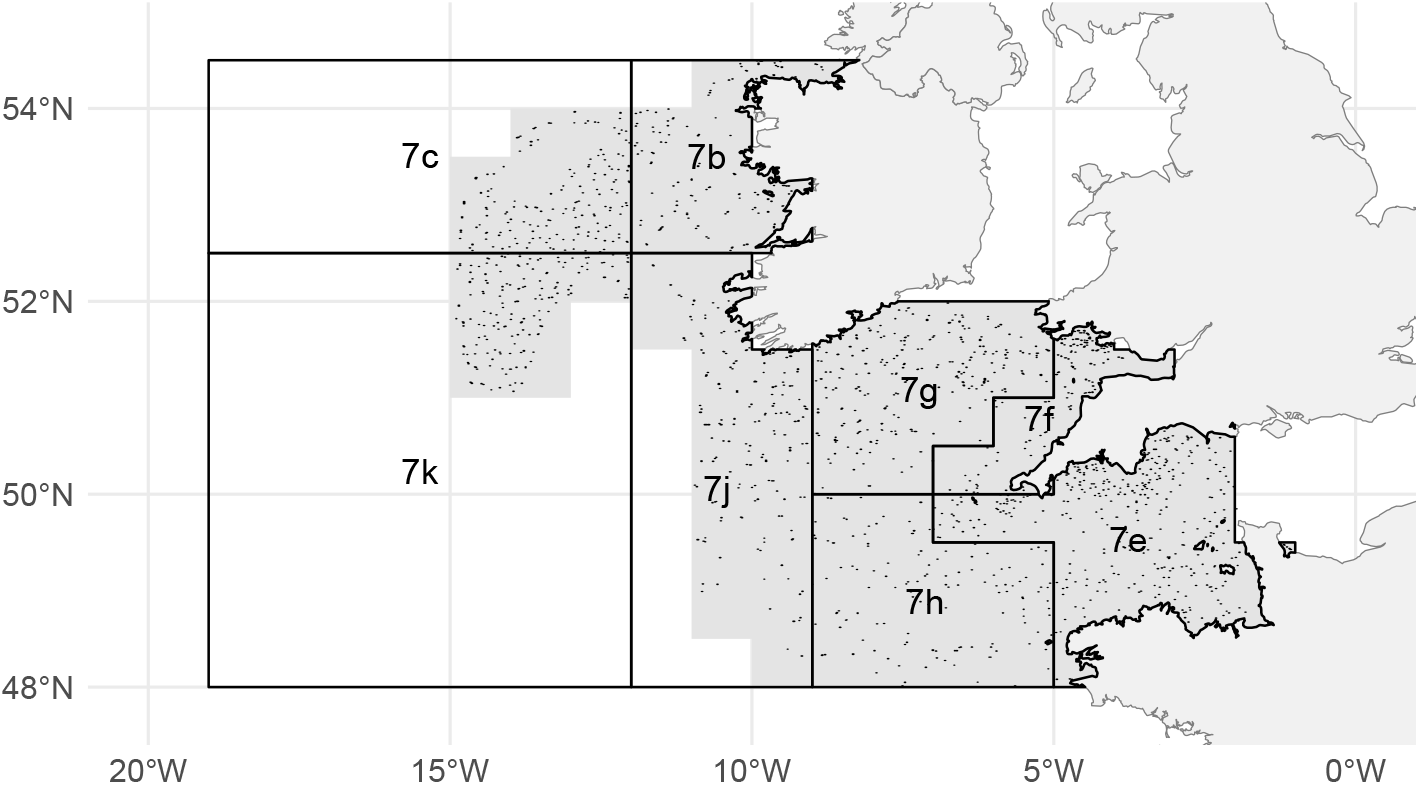
Map of ICES Subarea 7, excluding divisions a and d. The shaded area depicts the ICES rectangles surveyed, with the locations of 1876 survey hauls used in this study shown as points.

We investigated species ranked 1 to 67 by abundance, out of a total of 168 species caught in the surveys. These span >10000-fold range of population density, so rare as well as common species are well represented (Fig. 4). The species were mostly teleosts (Actinopteri), but also included 15 elasmobranchs and two rabbit-fishes (Holocephali). The two most common species, blue whiting *Micromesistius poutassou* (rank 1) and herring *Clupea harengus* (rank 2), are not strictly demersal, but are included in view of their abundance in the survey data. The ranked species list is in Table 1 (Appendix A); the method by which the densities were estimated is given in Appendix B; confidence intervals were obtained by bootstrapping at the level of hauls with spatial stratification in place (Appendix F).

**Table 1:**
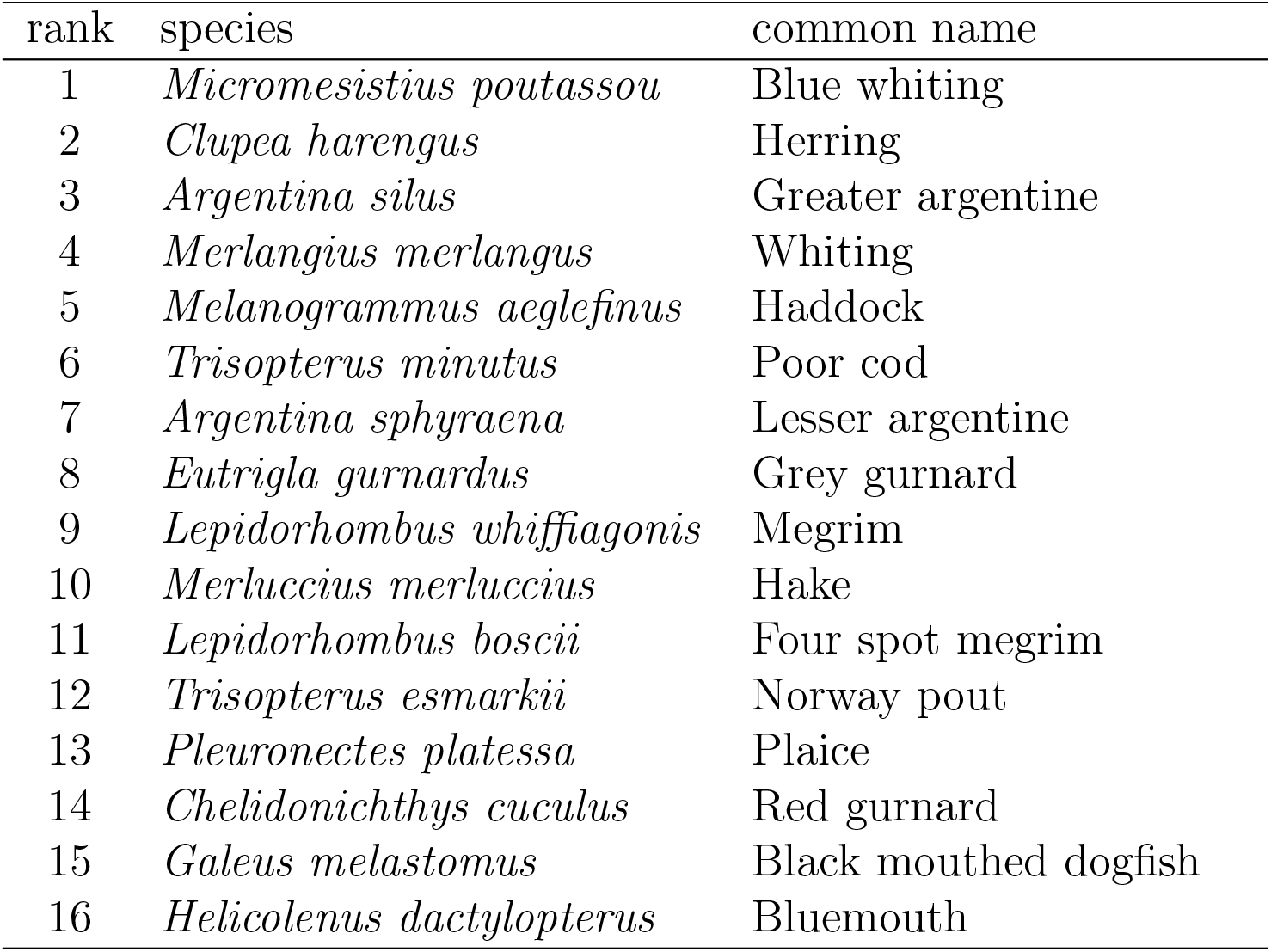
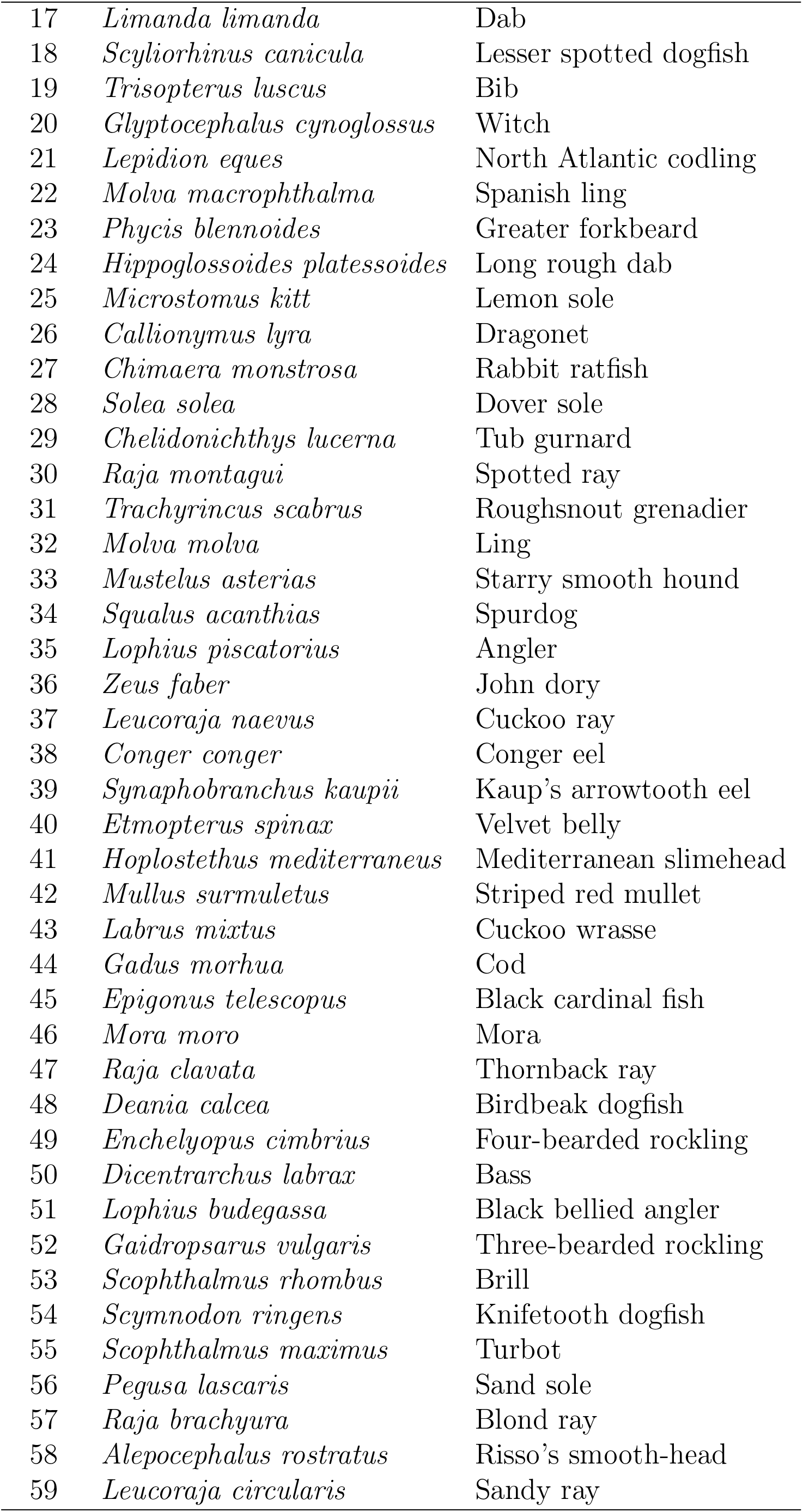
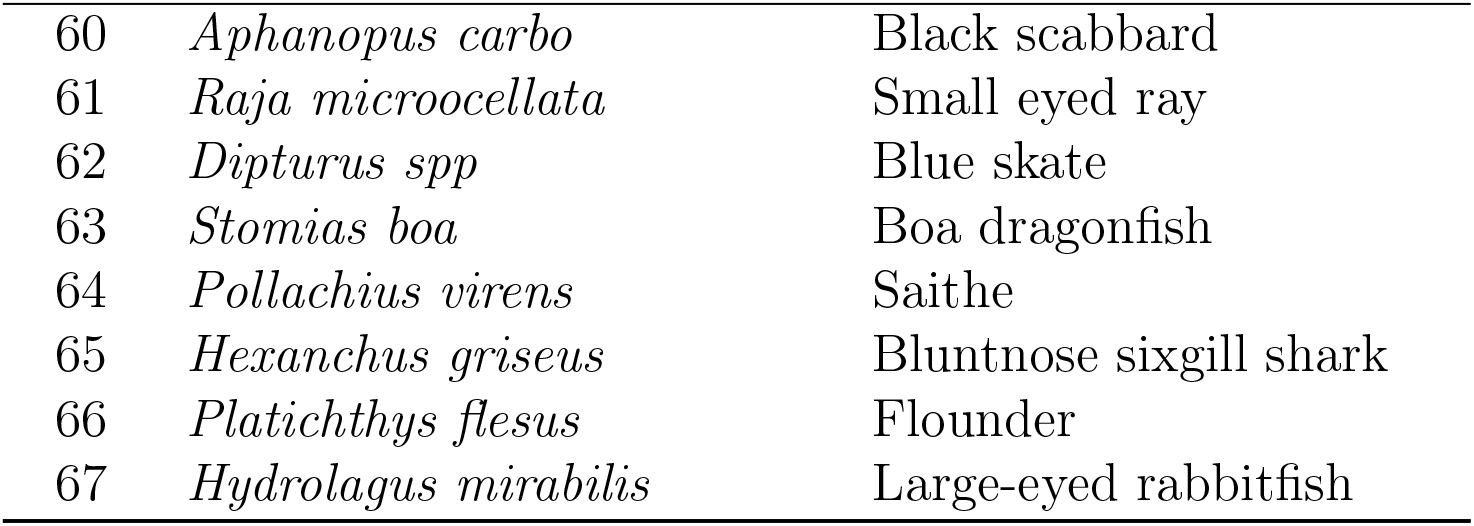
Species list ranked by density.

**Figure 4:**
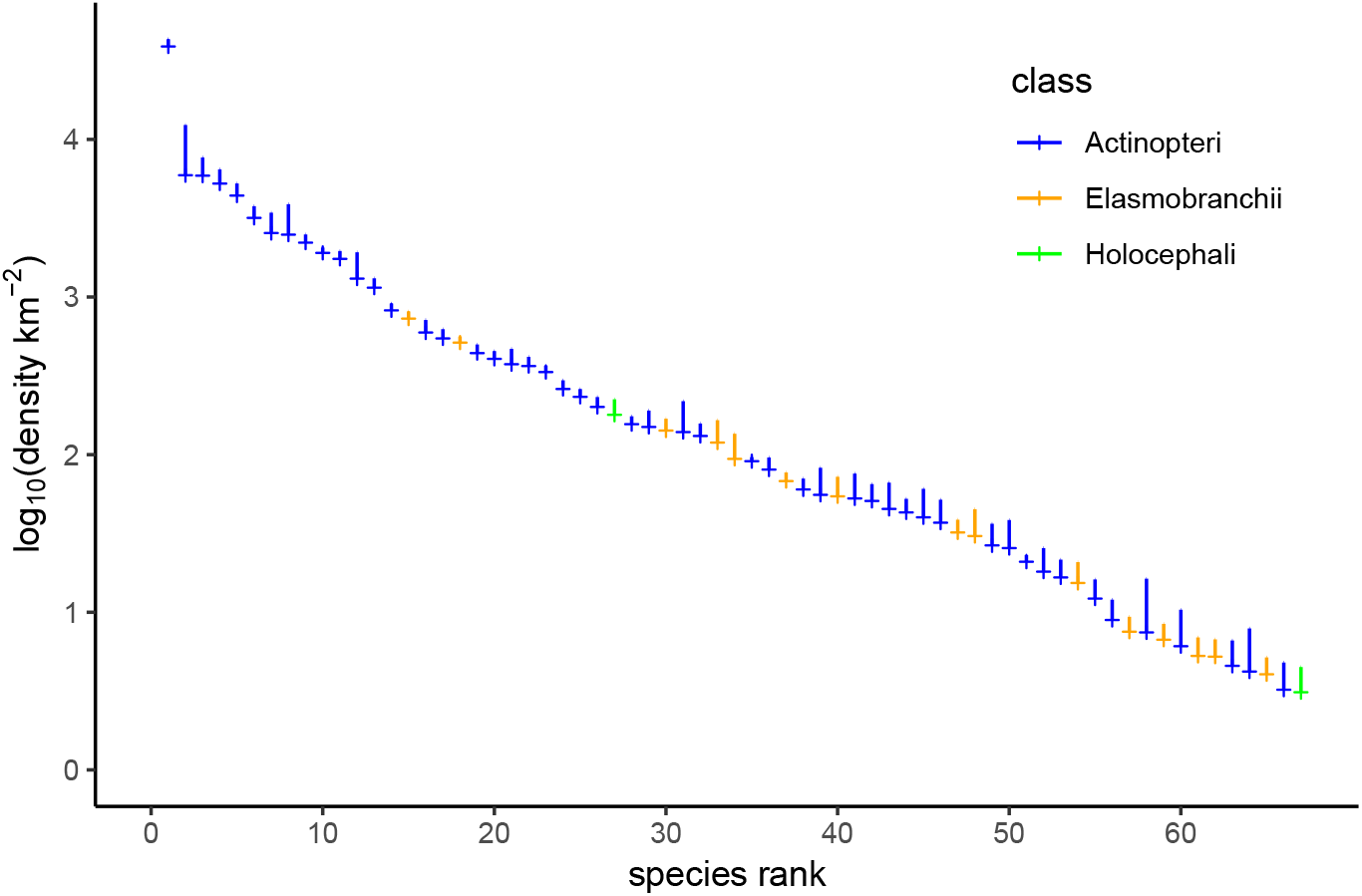
Species abundance distribution for demersal fish species in the Celtic Seas, estimated from otter- and beam-trawl survey data collected over the period 2012-16 from ICES divisions 7b,c,e-k. Mean densities with upper 95 % confidence limits are shown for 67 species recorded at least 100 times in the combined survey hauls, and with a minimum body length of 20 cm. See Table 1 in Appendix A for the ranked species list.

Fishing mortality rates *F*_*s*_ for each species *s* were estimated from fishing effort data available through EU STECF (Zanzi and Holmes, 2017). These data are classified *inter alia* by year, commercial fishing gear and ICES rectangle (rectangles of 30 min latitude by 1-degree longitude used for gridding of data). As a measure of effort, we took the ratio of the area swept yearly by mobile demersal fishing gear relative to rectangle area. This approach was adopted to allow comparability of the fishing mortality rates, as all species in the assemblage could then be treated in the same way. But it is important to be aware that static gears cannot be included in a swept-area ratio, so the *F*_*s*_s given here underestimate the fishing mortality. Also, some bias in sampling the fish assemblage by commercial vessels is to be expected as fishers seek profitable species to catch.

A fishing mortality rate was obtained for fish in each row of the survey data from the fishing effort, together with the catchability of the fish according to its species group, the kind of commercial gear and body length, as described in Walker et al. (2019) (see Appendix C). The fishing mortality rate *F*_*s*_ of species *s* was taken as the median value across all individuals of species *s* in the survey; confidence intervals were obtained by bootstrapping (see Appendix F). Fig. 5 shows that, although there was substantial variability in death rates from fishing, there is no indication that fishing mortality rates were lower in rarer species caught as bycatch, than in more common targetted ones. In other words, there is no evidence to suggest that rarer species get extra protection from fishing (see Section 2). It is notable that elasmobranchs, which account for 25 % of the taxa in the assemblage, make up 38% of the species for which *F*_*s*_ ≥ 0.05 yr^*−*1^.

**Figure 5:**
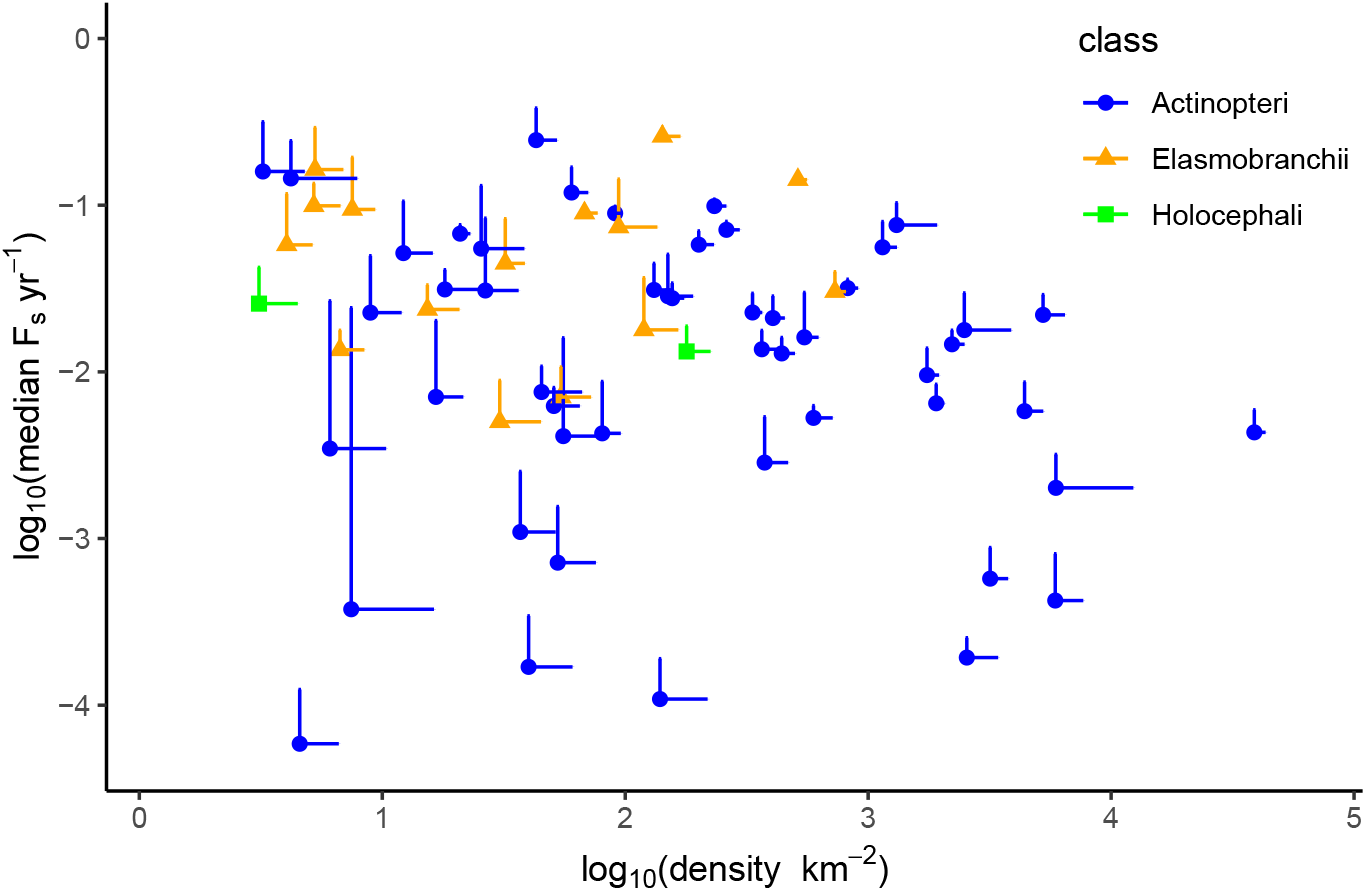
Absence of a relationship between median death rate from fishing (*F*_*s*_) and population density in 67 demersal fish species in the Celtic Seas, estimated from otter- and beam-trawl survey data collected over the period 2012-16 from ICES divisions 7b,c,e-k. Points are estimates for species with upper 95 % confidence limits shown as bars.

Fig. 6 steps from density to biomass density, and from fishing mortality to yield. Both steps introduce the mass of each surveyed fish. In the case of biomass density, this is evident from Appendix Eqs (B.1), (D.1), (D.2); in the case of yield, the harvest rate of each fish was multiplied by its body mass (Eqs (D.1) and (D.3)). Introducing biomass to both axes generates a relationship with slope ≈ 1, as would be expected in the absence of a relationship between fishing mortality rate and density in Fig. 5. The greatest yields, around 0.1 t km^*−*2^ yr^*−*1^, were from hake *Merluccius merluccius*, haddock *Melanogrammus aeglefinus* and whiting *Merlangius merlangus*. Note that ‘yield’ here refers to the rate of loss of biomass through commercial fishing at the point of capture, and is distinct from the rate at which biomass is landed in fishing ports.

**Figure 6:**
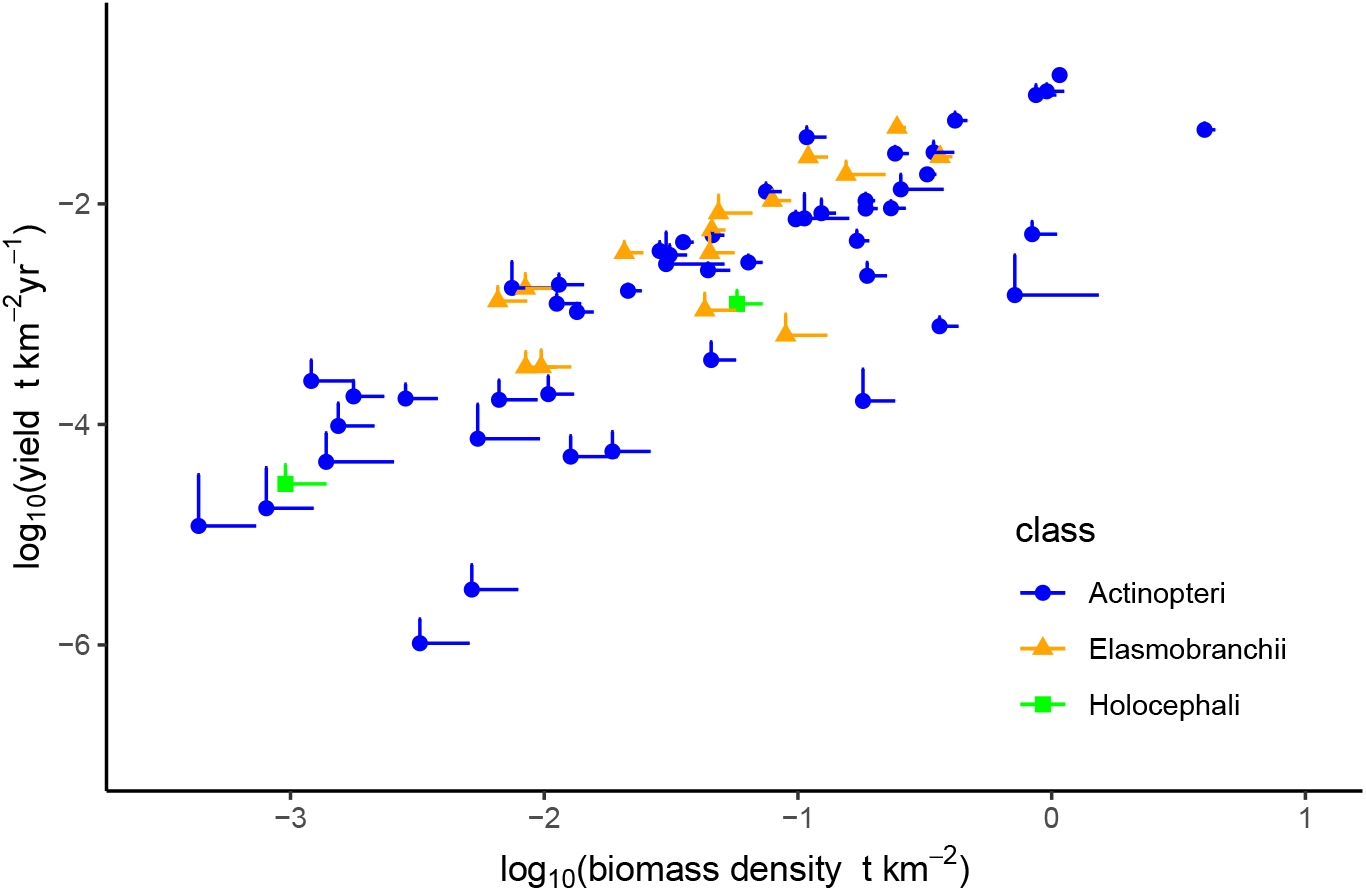
Rate of loss of biomass from fishing (yield) as a function of biomass density estimated from 67 demersal fish species in the Celtic Seas, estimated from otter- and beam-trawl survey data collected over the period 2012-16 from ICES divisions 7b,c,e-k. Points are estimates for species with upper 95 % confidence limits shown as bars.

We developed a measure of production rate motivated by dynamic size-spectrum models. Broadly, the contribution each fish makes to production at the time of sampling is the product of its mass and its mass-specific growth rate at that time (Morais and Bellwood, 2019). Summing this product over all individuals of a species, and dividing by the total area swept by the survey trawls, scales the production to a species-level measure with units t km^*−*2^ yr^*−*1^ (in the same way as the biomass from each surveyed fish scales up to a biomass density with units t km^*−*2^). Production rate measured in this way is conceptually distinct from that used in aggregated single-species, surplus-production models, where production is taken as an unstructured, logistic-like term (Pella and Tomlinson, 1969; Polacheck et al., 1993; Haddon, 2023).

In more detail, each surveyed fish was assigned a unique von Bertalanffy growth equation, with growth coefficient 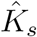, and asymptotic body length 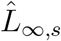 drawn from a bivariate normal distribution with mean ln*K*_*s*_, ln*L*_∞,*s*_ for each species *s*; *K*_*s*_, *L*_∞,*s*_ were obtained using functions in García-Carreras et al. (2016) and Walker et al. (2019). (The parameter *t*_0_ was assumed to be zero.) After transforming from length at capture to weight, the growth equation then gives the mass-specific growth rate for each fish at the time of capture (Eq. E.4), and hence its contribution to production of biomass. A bias is generated from taking growth as a function of body weight, rather than age, because slower growth reduces the chance of survival to a given size; we made a correction for this (Eq. E.7). A full explanation of the production calculation is given in Appendix E; confidence intervals were obtained by bootstrapping (Appendix F).

The relationship between log*Y* and log*P* (Fig. 7) has a linear regression slope 0.827 (SE: 0.110). This is consistent with the absence of a relationship between fishing mortality rates and abundance of species shown in Fig. 5. But importantly, it is not consistent with protection of rare species, according to dynamic models of multispecies assemblages. This theory suggests that rare species actually need more protection from fishing, i.e. lower fishing mortality rates, than common species. For instance, balanced harvesting would require a slope 1 +*α*, where *α* is the gradient of the relationship between log*P* and log*B* (Section 2), ≈ 0.865 in the Celtic Seas (Fig. 2). Balanced harvesting of this assemblage would therefore call for a log*Y*, log*P* relationship with a slope in the region of 1.865, also shown in Fig. 7. We caution against over-interpretation of this line, because the intercept log*c* in Eq. (2.9) is a free parameter that sets the overall intensity of fishing. Here log*c* = −1, a value chosen to bring the line near to the yields of three important commercial species, hake *Merluccius merluccius*, haddock *Melanogrammus aeglefinus* and whiting *Merlangius merlangus*.

**Figure 7:**
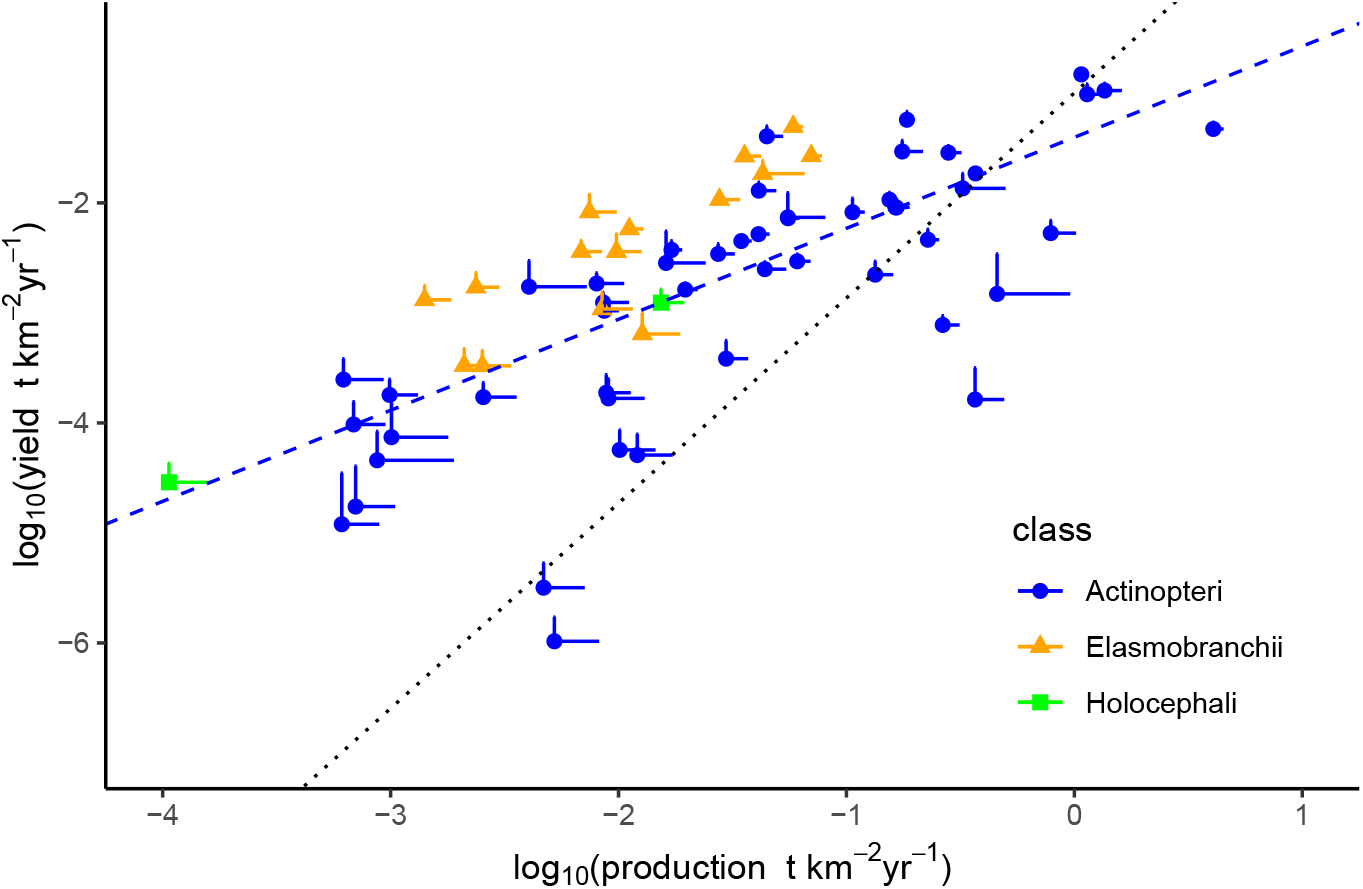
Rate of loss of biomass from fishing (yield) as a function of production rate estimated from otter- and beam-trawl survey data collected on 67 species in the Celtic Seas from ICES divisions 7b,c,e-k over the period 2012-16. Points are estimates for species with upper 95 % confidence limits shown as bars. The dashed line is a linear regression line through the points; the dotted line has a slope consistent with balanced harvesting. ‘Yield’ here is estimated at the point of capture, and is distinct from the landed catch.

We note that there is some separation of Elasmobranchii from Actinopteri in Fig. 7. For a given level of yield, species of Elasmobranchii tend to have lower production rates than species of Actinopteri. The reason for this is that production rate depends in part on the rate of body growth (Eq. E.9), and this is less in Elasmobranchii than in Actinopteri (see Eqs (E.2), (E.3), Appendix E). The low production rates of Elasmobranchii highlight the vulnerability of these taxa under contemporary patterns of exploitation.

## 4 Discussion

The evidence from this paper is that fishing mortality rates on rare species with low rates of biomass production were not lower than those on common species in the demersal fish assemblage of the Celtic Seas, during the period 2012 to 2016. The surveys used are the best information available, although there are caveats about inferring ‘rarity’ from them, discussed below. This result is not unexpected, because towed fishing gears in general use are largely non-selective. We think it is unlikely that the situation for rare vulnerable species has changed substantially since the time period 2012 to 2016. The main development has been the introduction of the Landing Obligation to prevent discarding (European Union, 2013), but this applies only to species of commercial interest regulated by total allowable catches (TACs) in the Celtic Seas, and not to rarer, unregulated species. Towed gears with some capacity for discrimination between species might help (Krag et al., 2015, 2016), but such gears are not in widespread use in the Celtic Seas.

The relationship between yield and production that emerges from the estimated fishing mortality rates is cause for concern. Results from dynamic models of fish assemblages suggest that species with low production rates need extra protection from fishing if they are to persist in exploited ecosystems (Law and Plank, 2023). But there is no sign of this protection in the results presented here, and this leads to the expectation that low-production species were likely to have been in decline. Consistent with this, empirical evidence from the Celtic Seas shows that species with large maximum body lengths (*L*_*max*_) were decreasing in abundance over the period 1986 to 2004 as a result of size-selective fishing (Shephard et al., 2012). The majority of species with *L*_*max*_ ≥ 75 cm in our study were Elasmobranchii (Fig. 8), facing dual threats from (i) low production rates, and (ii) high catchability due to large body size, which makes them especially vulnerable. Important also are smaller species with low production rates; these are easily forgotten, as they are unlikely to leave a signal in the Large Fish Index (LFI) (Greenstreet et al., 2011; Shephard et al., 2012).

**Figure 8:**
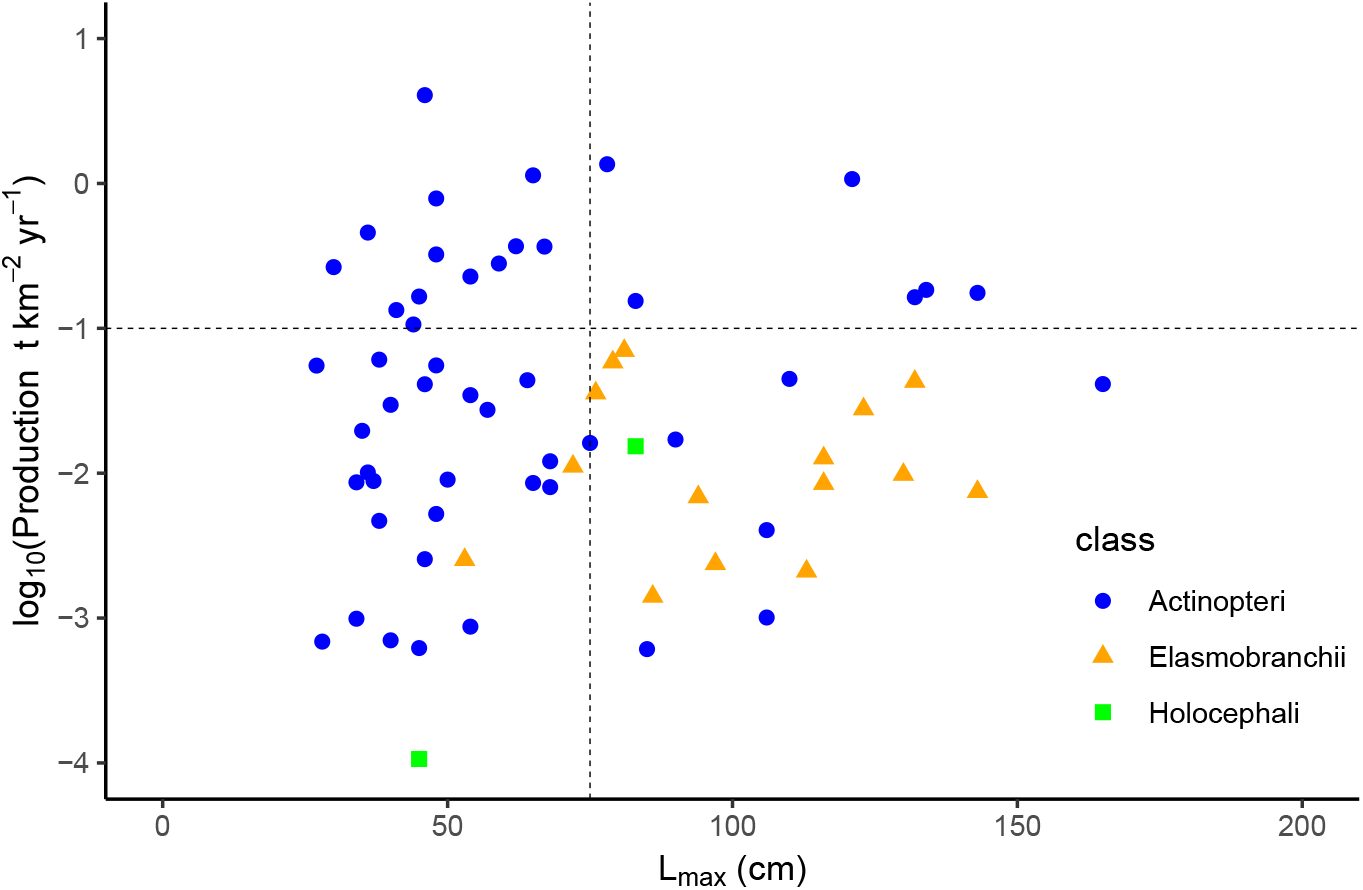
Relationship between production rate and maximum body length *L*_*max*_ in 67 demersal fish species in the Celtic Seas from ICES divisions 7b,c,e-k over the period 2012-16. *L*_*max*_ is taken as the largest individual observed in the survey data used in this study. Points are estimates for species.

We have no reason to believe that the Celtic Seas are exceptional among exploited marine ecosystems around the world. A meta-analysis of 110 Ecopath models from around the world found still shallower log*Y*, log*P* slopes than that in the Celtic Seas (Kolding et al., 2016), which implies fishing mortality rates that are greater on species with low production rates than on species with high production rates. This suggests the problem is widespread and deep-seated. The indications are that exploitation of marine ecosystems is not in line with Goal 14 of the United Nations Agenda for Sustainable Development.

To bring exploitation in marine ecosystems towards sustainability, the evidence from modelling multispecies assemblages is that fishing mortality rates would need to be moved in the direction of proportionality with production rates of species (Law and Plank, 2023). Such exploitation is known as balanced harvesting (Garcia et al., 2012; Heath et al., 2017; Zhou et al., 2019; Law and Plank, 2023). In practical terms, balanced harvesting calls for more protection to species with low production rates, relative to those with high production rates.

We note that some advice to fishery managers is moving in this direction. The ICES guidelines for a class of data-limited fish stocks contain a target reference length for the mean length at capture with the property *F* = *M* (ICES, 2024, Methods 2.1, 2.2, pages 15, 19). On its own, this would set the target reference length to an exploitation ratio *E* = 0.5, Eq. (2.3). However, if biomass is declining, the guidelines override this with a biomass safeguard, thereby reducing *F*. Bearing in mind that biomass is a key component of production rate, the biomass safeguard can be interpreted as a step towards aligning fishing mortality with production rate. The work here suggests a further step of embedding biomass directly into the production rate, this being the natural counterpart to the rate at which biomass is removed as yield. Our work shows that this step can be made using standard fisheries information, which means that the sustainability of exploited fish assemblages can then be assessed through log*Y*, log*P* plots. The approach taken here is a first step in this direction, and there is plenty of scope to develop it further.

The total rate of loss of biomass from fishing (‘yield’) estimated from the survey was approximately 1.3 t km^*−*2^ yr^*−*1^ (Appendix D). This is comparable with total yields from the Celtic-Biscay large marine ecosystem (LME) estimated in the range 1.5 to 2 t km^*−*2^ yr^*−*1^ over the same time period (Ryther index, Link and Watson, 2019), and a maximum multispecies sustainable yield (MMSY) of 1.8 t km^*−*2^ yr^*−*1^, computed from a dynamic size-spectrum model (Jennings and Collingridge, 2015, Fig. 8, scenario A). The lower value from the survey would be expected, because ICES Subarea 7 extends further west from the shelf than the LME, although there are also losses between the point of capture in the surveys and the point of landing that need to be taken into account. However, these values are all substantially greater than the total yield given by the EU STECF landings data, which was 0.4 t km^*−*2^ yr^*−*1^, in ICES divisions 7b, c, e-k over the time period 2012 to 2016. Taken at face value, this would imply less than a third of the total catch was landed. This is unlikely, and we have not been able to find an explanation for the discrepancy.

Log*Y*, log*P* plots hold some promise as a measure of health of exploited marine ecosystems (Jennings, 2005; Heath et al., 2017; Fulton et al., 2025). They provide an overview of the relative levels of exploitation of an assemblage of species, encompassing rare as well as common ones, and are underpinned by a theoretical framework that describes dynamics of multiple, interacting species (as opposed to single, isolated species). Moving fishing mortalities in the direction of alignment with production rates of species can be thought of as an ecosystem harvest control rule (EHCR) (Garcia et al., 2016) that protects vulnerable species and may leave room for greater exploitation of certain common species, if their production is shown to be underutilised. It raises a possible certification of fisheries at an ecosystem level, without incurring the costs of single-species certification. Overall, the EHCR brings the agendas of fisheries exploitation and conservation together on a common platform. Note however, that the focus of this study is on the *relative* levels of exploitation of species within an ecosystem, not the total rate at which the ecosystem is exploited. The latter is a societal matter involving many stakeholders, and is ultimately constrained by the rate of primary production (Chassot et al., 2010; Link and Watson, 2019): balanced harvesting would argue for the need to keep exploitation at a moderate level (Garcia et al., 2012).

General conclusions from these results on the Celtic Seas should be made with several provisos. Strictly speaking, the results apply only to the 96 ICES rectangles surveyed in the Celtic Seas, as we have no information on the other 85 ICES rectangles. The death rates from fishing, being based on towed gears, underestimate the *F* s for species that are also caught in static gears. Estimates of abundance based on demersal trawls also miss the important guild of pelagic species. We did include two pelagic species, herring (*Clupea harengus*) and blue whiting (*Micromesistius poutassou*) as these were among the most common species in the surveys, but pelagic surveys could give still greater values for their abundance. There is bound to be some mismatch between the assemblage as sampled by surveys, and that sampled by fishers as they seek out locations where their target species live; this also contributes to underestimation of *F* s of commercially-important species in our study. It is important to keep in mind that the demersal fish assemblage is just a small part of the biodiversity of Celtic Sea ecosystems. For instance, most of the pelagic species of Actinopteri were not included, and invertebrate species, some of which are commercially important, were not taken into account. Also excluded were the plankton, key components of marine ecosystems, as were marine mammals and birds which are of great importance for conservation (Cury et al., 2011).

It is also important to bear in mind that species may be caught rarely in the survey gear for reasons other than inherent rarity. For example, Flounder (*Platichthys flesus*), which has a coastal distribution, is rarely caught in the surveys (Lart, 1986). A species may appear to be rare because it is passing through the surveyed area on migration. A species could be close to its maximum length at the minimum body length of 20 cm used in our study; while rare at this length, it would be more common at smaller sizes. We excluded Imperial scaldfish (*Arnoglossus imperialis*) on these grounds, as it had a maximum body length of 23 cm. Clearly, many species with maximum body lengths < 20 cm are excluded altogether, including Boarfish (*Capros aper*) and mesopelagic taxa.

Despite these limitations, this study shows that production rates can be estimated systematically in multispecies assemblages, using information standardly collected for fisheries management. Production, taken here as the rate of accumulation of biomass, is the direct counterpart of its rate of removal as yield or bycatch, and allows the relationship between production and exploitation to be examined in multispecies assemblages. The results suggest that conservation of rare species, which are not themselves targets of fishing, is not achieved by current methods of exploitation. In working towards the UN goal of sustainable development (SDG14), production rates of both common and rare species need to be monitored, and mortality rates from fishing adjusted to bring them towards a balance with their rates of production.

## Acknowledgements

This work was supported by UK Research and Innovation’s Sustainable Management of Marine Resources programme, the Pyramids of Life: Working with Nature for a Sustainable Future project [grant number NE/V016008/1]. We thank Gustav Delius, William Lart and the Pyramids of Life team for their input into the work.

## APPENDICES

### A Survey data

The data for the Celtic-Sea fish assemblage used in this study came from the International Bottom Trawl Survey (IBTS) which uses a standard otter trawl ‘Grande Overture Vertical’ (GOV) (ICES, 2017), and the Beam Trawl Survey (BTS) (ICES, 2009). The study was intended as a spot check on the assemblage rather than as an analysis of changes over time, and therefore did not require a long time-window. But the window still needed to be long enough to achieve adequate samples of at least some of the rarer species. So a five-year period from 2012 to 2016 was chosen as the most recent time interval for which complete fishing effort and landings data are publicly available from the European Union Scientific, Technical and Economic Committee for Fisheries (EU STECF). The surveys took place in 96 out of the 181 ICES rectangles in Subarea 7 Divisions b, c, e-k during the period of study, about 50 % of the area. However, 98 % of the commercial fishing effort (of the gears used here), took place in the surveyed rectangles during this time (EU STECF effort data), so the spatial match between the surveys and the commercially fished areas was close. Almost all surveys near SW England used beam trawls (Divisions 7e, f); west of this is a region where both beam and otter trawls were used (Divisions 7g, h); west of this and west of Ireland almost all surveys used otter trawls (Divisions 7b, c, j, k). The study was based on a total of 1876 hauls (GOV: 1174, BTS: 702), 1872 of which contained fish used in this paper. Seasonal changes in the fish assemblage are not considered.

Typically, a single row of data in the survey files corresponds to a single captured fish, including information on: location and date of capture, species and body length of the fish, a haul identifier and the area swept by the haul. Our calculations added to this: (i) a fishing mortality rate (yr^*−*1^), (ii) the contribution to yield (t km^*−*2^ yr^*−*1^), and (iii) the contribution to production rate (t km^*−*2^ yr^*−*1^). However, some rows contained multiple individuals, if a large number of fish of a particular species and body length were caught in a haul and then subsampled. In such cases, the number of fish in a row at the time of capture was estimated from the total weight divided by the weight of a subsample with a known number of individuals. We write the number of individuals in row *i* of the survey data as *n*_*i*_.

Following the advice of Walker et al. (2019), the study was based on fish ≥ 20 cm in length. A species could appear to be rare because this lower limit is close to the maximum size to which individuals grow. Imperial scaldfish *(Arnoglossus imperialis)*, with a maximum observed body length of 23 cm, and only 142 individuals in the surveys in the length range 20 to 23 cm, was excluded on these grounds. Pelagic species were excluded because the survey gears used in the IBTS and BTS surveys are unsuitable for sampling them. This is with the exceptions of Blue whiting (*Micromesistius poutassou*) and Herring (*Clupea harengus*), which were among the most common species in the surveys. The study was restricted to the 67 species listed in Table 1 (out of 168 species all told) that were sampled at least 100 times in the combined data files. Skate species were aggregated into a single group listed as *Dipturus spp*. After filtering, the study was based on a total sample of ~2.4 million fish.

### B Species abundance distribution

To get estimates of the abundances of the species in Table 1, requires their catchabilities in the survey gears, which are themselves dependent on body size. It would have been difficult to get this information separately for each species, so we adopted a method used in the North Sea by Walker et al. (2017), assigning each species to one of seven species groups according to body shape and position in the water column of the species (Walker et al., 2017; Rindorf et al., 2020):

1. predominantly buried in sediment
2. on or near the seabed – anguilliform or fusiform
3. predominantly on the seabed – flat
4. predominantly close to the seabed, but not on it
5. midwater species with some seabed association
6. pelagic
7. predominantly on the seabed – lumpiform.

In most cases species groups were available from the literature (Walker et al., 2017; Rindorf et al., 2020); Fishbase was used in a small number of cases (Froese and Pauly, 2000).

Walker et al. (2017) provided catchability, 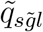, disaggregated by survey gears 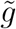 (IBTS, BTS), body length *l*, and species *s* (simplified here to species group). (The tilde denotes ‘survey gear’, which needs to be distinguished from commercial gears below.) We assigned a catchability to each row *i* in the survey data as 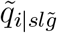, according to *s, l*, 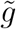 in row *i* of the survey data. The estimated number of fish in the sea generating the catch in row *i* of the survey data is then 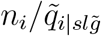, where *n*_*i*_ is the number of fish in row *i* (Appendix A). The total number of fish generating the catch of species *s* is the sum of this ratio over gears and body lengths. From this an estimate of population density, *X*_*s*_ (km^*−*2^), of species *s* in the sea, can be obtained from the area swept by all survey trawls *Ã* as:

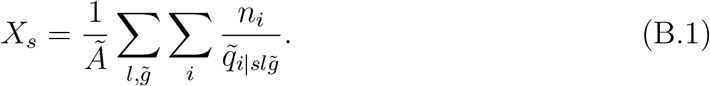

Population densities of the species obtained in this way are displayed as a species abundance distribution in Fig. 4.

### C Death rate from fishing

The aim was to attach a fishing mortality rate to every row *i* in the survey data, according to the species, body length, ICES rectangle and year in row *i*. We adapted a scheme used in the North Sea by Walker et al. (2019) to do this, combining records on fishing effort with catchability of commercial fishing gears.

Fishing effort was obtained from a data file ‘effort-by-ICES-rectangles.xlsx’ (downloaded: 02.10.23) from the EU STECF (Zanzi and Holmes, 2017), extracting from this the data sheet CEL1 which covers ICES Subarea 7b,c,e-k. To match the area of the survey data, the sheet was filtered down to the 96 rectangles used in the survey, and the time period 2012 to 2016. English and Scottish efforts had been given to EU STECF as ‘number of days at sea times 24’ rather than as the number of hours of fishing, and were corrected using the factors given in Engelhard et al. (2015, Table S1); the effect of this correction is appreciable.

We took the swept area ratio as a measure of effort, using the yearly total area swept by fishing gear in each rectangle divided by the rectangle area. (The yearly time-frame gives units yr^*−*1^ to the death rate from fishing.) Since the data file gives effort in hours rather than area fished, Table 1 of Walker et al. (2019) was used to make a transformation to a gear-dependent swept area. Basing fishing effort on the swept area ratio means that the study was restricted to fishing mortality from mobile gears: inclusion of static gears would be needed to get the total death rate from fishing. The study was confined to commercial gears comprising otter trawls (coded TR1, TR2) and beam trawls (coded BT1, BT2), as these accounted for 97 % of the effort from mobile gears in the surveyed rectangles over the period 2012 to 2016. The output from this filtering was a measure of fishing effort *e*_*ryg*_, where *r* is an index for ICES rectangle (as in the survey data), *y* is an index for year (*y* = 2012, …, 2016), and *g* is an index for commercial gears TR1, TR2, BT1, BT2, after summing over contributions by country, vessel length class and quarter.

Conversion from effort to fishing mortality needs catchability for the four mobile commercial fishing gears (TR1, TR2, BT1, BT2), resolved to species group and body length. This was taken from a data file ‘efficiencytable.csv’ in Walker et al. (2017). We denote this as *q*_*slg*_, indexed by species *s*, body length *l* and gear *g*, bearing in mind each species was assigned to a species group (Appendix B). The product of catchability and effort gives a death rate from fishing *f*_*slryg*_ = *q*_*slg*_*e*_*ryg*_. Since the study is concerned with the death rate aggregated over fishing gears, the fishing mortality rate *f*_*slry*_ is taken as:

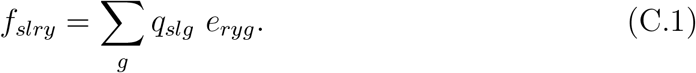

This death rate from fishing can then be assigned without ambiguity to each row *i* of the survey data as *f*_*i*|*slry*_, given the species *s*, body length *l*, rectangle *r* and year *y* of the fish recorded in row *i*. (We adopt a notation here and below of a lower-case variable name for measures recorded at the level of rows in the survey data file, and an upper-case name for measures aggregated to a species level; for fishing mortality, these are *f* and *F*.)

As a summary statistic for species *s*, the median value of *f*_*i*|*s*_ in all rows of the data containing species *s, F*_*s*_, is used in Fig. 5. This is preferred to a mean value because the body-length distributions tend to be skewed with more smaller than larger individuals, and the median is less affected by this asymmetry. In calculating the median value, we allowed for the catchability of the fish in each row *i* of the survey gear, disaggregating row *i* to 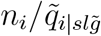.

### D Biomass and yield

The contribution of row *i* to the biomass of species *s* at length *l* caught in survey gear 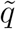 was estimated as

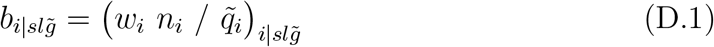

where *w*_*i*_ is the mass of a fish in row *i*, obtained from its body length *l*_*i*_ as 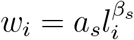, with allometric parameters *a*_*s*_, *β*_*s*_ for species *s* given in the survey data files. Biomass density of species *s* (units: t km^*−*2^) is then estimated as

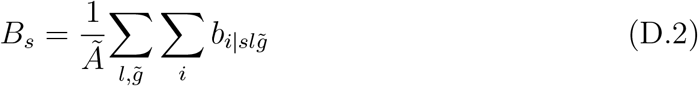

where *Ã* is the total area covered by the survey trawls. Values of *B*_*s*_ estimated in this way are used in Fig. 2.

The contribution of row *i* of the survey data to the yield uses the harvest rate *h*_*i*|*slry*_, matched to the rectangle *r* and year *y* of row *i*. This is a function of the fishing mortality rate *f*_*i*|*slry*_ (Eq. (C.1)), *h*_*i*|*slry*_ = (1 − exp(*τf*_*i*|*slry*_))*/τ*, with a time constant *τ* = 1 year for dimensional consistency. The product *h*_*i*|*slry*_ with 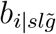 (Eq. (D.1)) is the contribution of row *i* to the yield of species *s* at length *l*. The overall yield density *Y*_*s*_ from species *s* (t km^*−*2^ yr^*−*1^) is then the summation of the product:

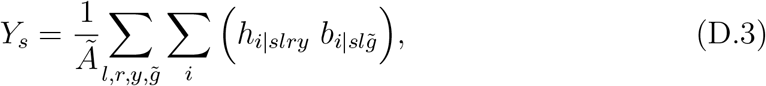

Values of *Y*_*s*_ estimated in this way are used in Figs 6 and 7. Yield here is measured at the point of capture, and should be distinguished from a landed catch: it is just the first step in a sequence of events that lead eventually to landings of fish at a port.

We compared our yield at the point of capture with the yield measured at the time of landing using a landings data file ‘landings-by-ICES-rectangles.xlsx’ (downloaded: 02.10.23) from the EU STECF (Zanzi and Holmes, 2017), extracting from this the data sheet CEL1 which covers ICES divisions 7b,c,e-k. To match the survey data, the sheet was filtered down to the 96 rectangles used in the survey, the gears in the survey (TR1, TR2, BT1, BT2), and the time period 2012 to 2016. There were 35 species in the landings data in common with species in the survey data, and a rate of loss of biomass in the survey data from fishing on these species was 0.73 t km^*−*2^ yr^*−*1^. But these 35 species accounted for less than a third of the landings, so we raised our value by the ratio of the total landings to the landings of these 35 species, a factor 3.67. The surveyed rectangles account for about half the area of ICES subarea 7, and about 94% of the fishing effort. Assuming the landings matched the effort, that would bring the rate of loss of biomass to about 1.3 t km^*−*2^ yr^*−*1^ according to the survey data. A caveat is that, although the fishing effort outside the surveyed areas was small, it led to substantial landings, especially of Blue whiting *Micromesistius poutassou*, that would increase the rate of loss of biomass. We note a discrepancy between our estimate from the survey and the value 0.4 t km^*−*2^ yr^*−*1^ given by the landings in ICES Subarea 7 over the same time period, which we cannot account for.

### E Production rate

We describe here how the contribution of each surveyed fish to production rate of the assemblage was estimated. Production rate of a single fish was measured as the rate at which it was accumulating mass at the time of capture in the surveys, i.e. the product of its mass and its growth rate per unit mass taken from the von Bertalanffy growth equation. From this information on each surveyed fish, an aggregated measure of production rate *P*_*s*_ by each species *s* could be calculated. Other aggregations are also possible, such as by body length.

To get the von Bertalanffy growth coefficient *K*_*s*_, and asymptotic body length *L*_∞,*s*_ for a species *s*, we followed a method used by García-Carreras et al. (2016) and Walker et al. (2019). This starts from the length *L*_*max,s*_ of the largest fish of species *s* sampled, this being the basic piece of information available from the surveys on species’ life histories. The scheme then makes use of a relationship between *L*_*max,s*_ and *L*_∞,*s*_:

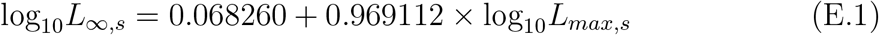

originally from Froese and Binohlan (2000), and updated by García-Carreras et al. (2016). There is evidence that *K* tends to be lower in species with larger *L*_∞_, and that overall the Elasmobranchii grow more slowly than the Actinopteri. We therefore used equations given by (García-Carreras et al., 2016; Walker et al., 2019) to describe and distinguish between the *K, L*_*oo*_ relationships of these groups:

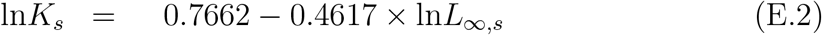

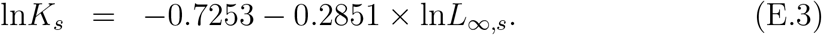

where Eq. (E.2) applies to Actinopteri and Eq. (E.3) to Elasmobranchii. Two species of the subclass Holocephali were present in the assemblage, and these are treated as Elasmobranchii. On completion of this step, each species *s* had been allocated parameters (ln*K*_*s*_, ln*L*_∞,*s*_) for the von Bertalanffy growth equation; the theoretical age at which length is zero, *t*_0_ was held at 0 throughout.

**Figure E1:**
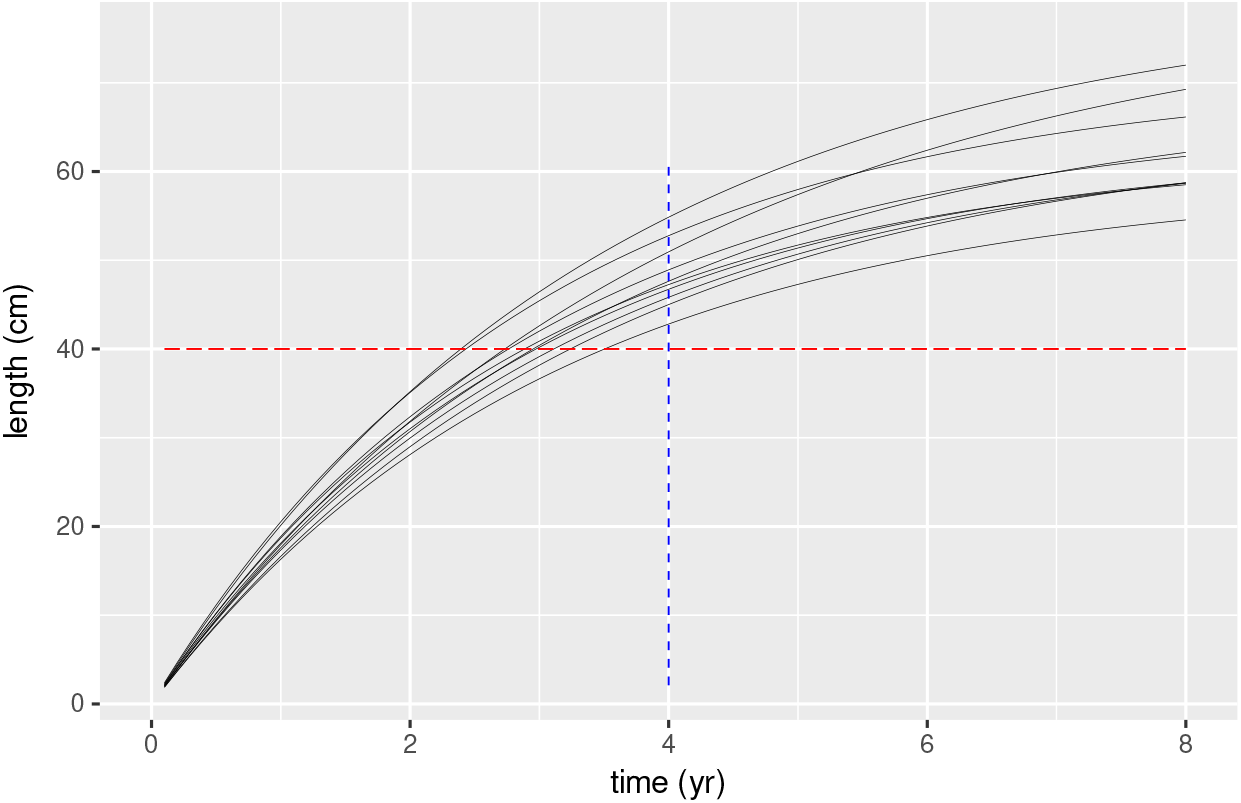
10 random growth trajectories of Whiting (*Merlangius merlangus*). Growth rate at a given length (40 cm) is the slope of the trajectory as it passes through the the horizontal line. Growth rate at a given age (4 years) is the slope of the trajectory as it passes through the vertical line.

Working at the level of individual fish in the surveys allows each fish to be allocated its own growth trajectory. Because some rows of the survey data contain multiple fish of a given species and body length, we started by disaggregating the survey data so that each row would correspond to a single fish caught. We then took the parameters for each species *s* (ln*K*_*s*_, ln*L*_∞,*s*_) as mean values for bivariate normal distributions from which to draw random pairs 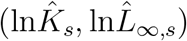, giving each individual its own growth trajectory. This needs a covariance matrix: the covariances were based on the regression parameters of Eqns (E.2), (E.3), and variances set 0.01 were found to give reasonable ranges for growth trajectories of individual fish. Thus the growth trajectory of a single fish of species *s* is taken as a von Bertalanffy growth equation with a parameter pair 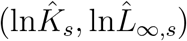, drawn at random from a bivariate normal distribution with means 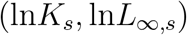 and covariance matrix **Σ**_*A*_ for Actinopteri, and **Σ**_*E*_ for Elasmobranchii. On completion of this step, each row *i* of the survey data had a unique growth trajectory. Fig. E1 gives an example of ten random trajectories of Whiting (*Merlangius merlangus*) obtained in this way.

As a preliminary to allocating a growth rate to each fish *i*, its length *l*_*i*_ at the point of capture was transformed to weight 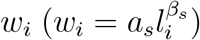, using species allometric parameters *a*_*s*_, *β*_*s*_ given in the survey data. The mass-specific growth rate of fish *i* with body weight *w*_*i*_ at the point of capture was then obtained from the time derivative of the growth equation evaluated at *w*_*i*_:

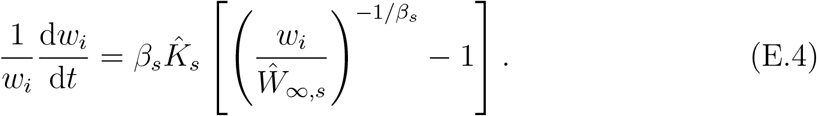

The geometry of this equation can be seen in Fig. E1 for fish caught at length 40 cm: their growth rates 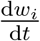 at this length are the slopes of the growth trajectories as they pass through the horizontal line at 40 cm (after transforming from length to weight).

It is important to keep in mind that fish with faster growth reach a given length at an earlier age than those with slower growth, and are likely to have accumulated a lower risk of death. So the growth rates at a given body length (e.g. 40 cm in Fig. E1) should be weighted by survivorship to this body length. There is no direct information on age in the survey data, but the von Bertalanffy growth equation can be re-arranged to give the time taken to grow from one length to another, so survivorship associated with growth at different rates can be taken into account. The time *τ*_1_ taken for a fish of species *s* to grow from length *l*_1_ to *l*_2_ is:

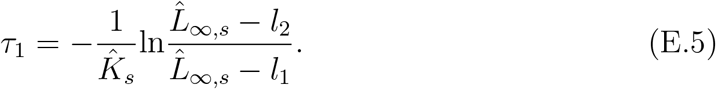

A length interval *l*_*min*_ to *l*_*max*_ can be divided into an arbitrary number of length steps *j* = 1, 2, …, with corresponding time intervals (*τ*_1_, *τ*_2_, …), and instantaneous rates of natural and fishing mortality (*m*_1_, *m*_2_, …), and (*f*_1_, *f*_2_, …) respectively. Survivorship *σ* through the full length interval is then

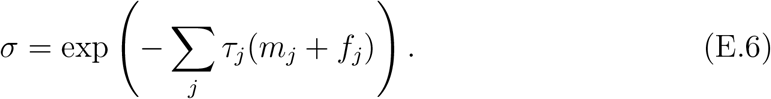

Eq. E.6 would have gone beyond the bounds of computability, so we took a simplified version, dividing growth into two stages of duration *τ*_1_, *τ*_2_, from 1 cm to 19 cm, and from 20 cm to length *l* at capture respectively. We assumed a fixed rate of natural mortality *m* = 0.2. yr^*−*1^ operated throughout this period, and a fishing mortality *f*_*i*_ already computed for each fish *i*, from 20 cm onwards. Survivorship of individual *i* of species *s* from 1 cm to its length *l* at capture is then

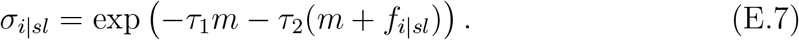

This oversimplifies the distribution of size-dependent fishing mortality as it does not allow for changes in catchability in the commercial gear before being caught in the survey. We checked that the results on production rate were not unduly sensitive to the assumptions made, but it would clearly be possible to take a much more nuanced approach to the mortality accumulated during growth.

Selecting rows of data from the surveys corresponding to species *s* caught at length *l* (*i*|*sl*), the survivorship weight for growth rate in row *i* is

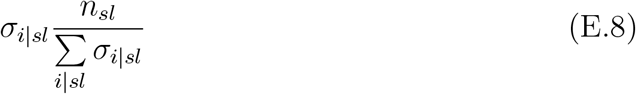

where *n*_*sl*_ is the number of fish of species *s* caught at length *l*. The contribution of row *i* to the production rate, given that row *i* corresponds to species *s* with length at capture *l*, is the product of the expressions in Eqs (D.1), (E.4) and (E.8)

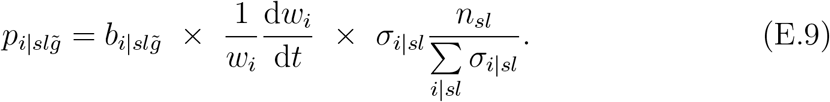

Note that catchability of the survey gears 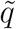 is accounted for in 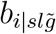 Eq. (D.1), so the contribution to production by row *i* refers to the population in the sea from which *i* was drawn, rather than the subpopulation caught in the survey nets. The total production rate *P*_*s*_ by species *s*, expressed per unit area, is:

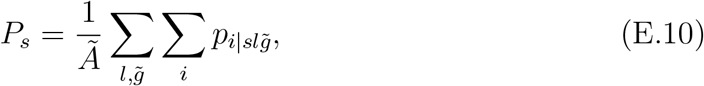

with units t km^*−*2^ yr^*−*1^. *P*_*s*_ is the species production rate used in Fig. 7.

### F Error estimates from bootstrapping

We used bootstrapping to obtain estimates of the errors associated with the aggregated measures for each species *s* from the surveys: *x*_*s*_, *B*_*s*_, *F*_*s*_, *Y*_*s*_, *P*_*s*_ **(ref?)**. This was done by sampling at random (with replacement) the 1872 hauls taken during the period 2012 to 2016 that contained the species and body sizes used in this study. Before doing this, we checked how much variation could be removed by a spatial stratification, comparing unstratified sampling (1 stratum) against: 8 strata (ICES divisions), 51 strata (pairs of ICES rectangles), 96 strata (ICES rectangles), taking 500 bootstrap samples in each case. The tests were carried out on species population density (*x*_*s*_) as a key measure in the study. The results showed the spatial stratification reduced the coefficient of variation of population density in 52 species (ICES divisions), 60 species (pairs of ICES rectangles), and 62 species (ICES rectangles). We chose the intermediate level of stratification (Table 2) on the grounds that this retained a substantial number of hauls within strata, while reducing the variability in about 90 % of the species (the reduction was often small). A stratified random sample structured as in Table 2 then entailed 21 hauls taken at random (with replacement) from stratum 1, 28 hauls from stratum 2, and so on, up to stratum 51. In this way, the number of hauls from a stratum remained as in the survey. The number of hauls was 1872, slightly reduced from the original number (1876) after filtering down to 67 species for the study.

**Table 2:**
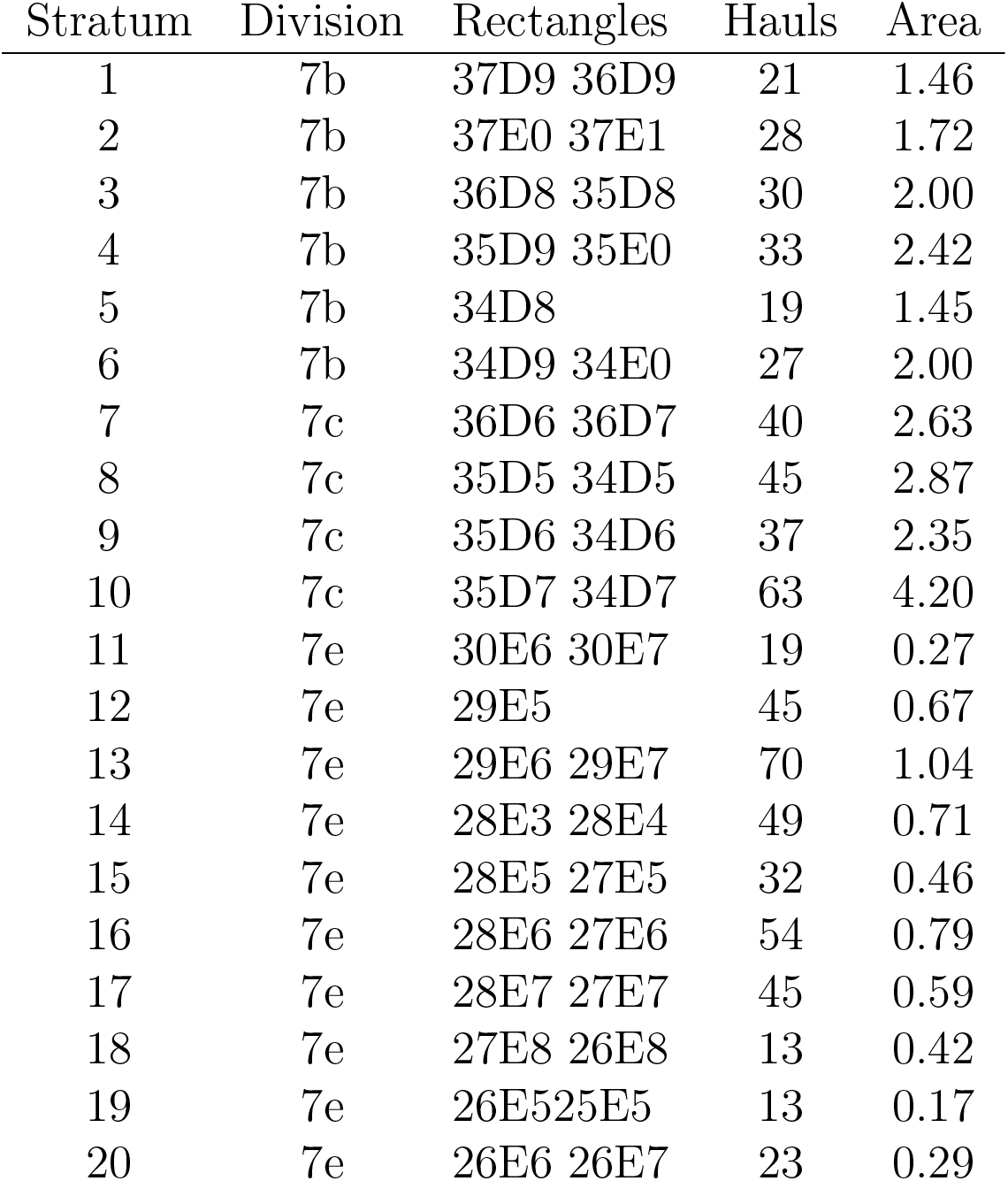
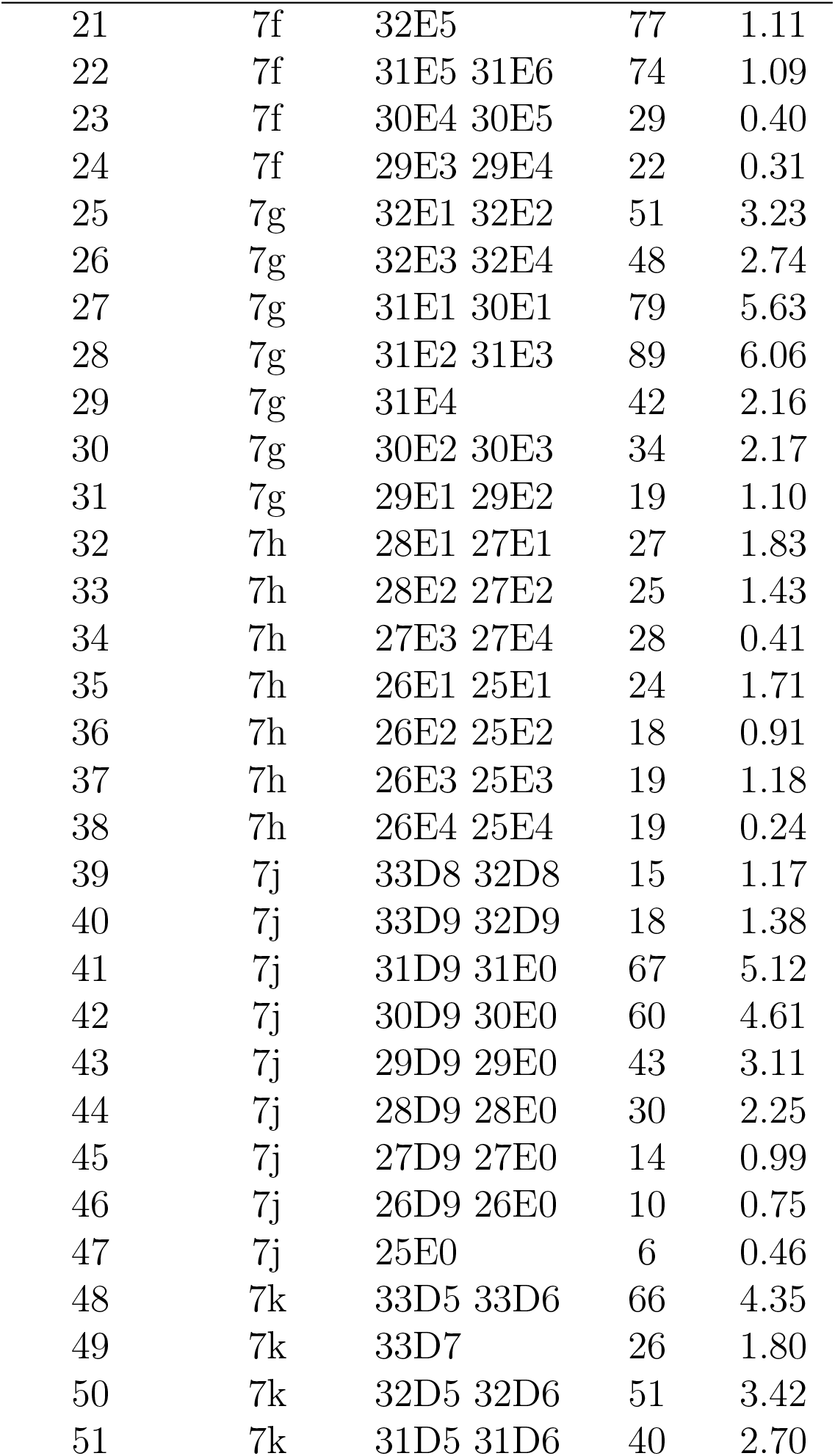
Stratification for bootstrapping, showing for each stratum: ICES Division, ICES rectangles, the number of survey hauls in the period 2012 to 2016, and the total area sampled by the surveys (km^2^) during this period.

Variation between bootstrap iterations is expected for many reasons, including for instance the sampling process itself, changes in abundance over the period of the survey, movement of species across the boundaries of strata, differences in the gear used by countries contributing the survey, differences between the beam- and GOV-trawls, and departures in catchability from the modelled values. We noticed a pattern of relatively low variation in bottom-dwelling Actinoperi, consistent with a relatively sedentary lifestyle.

In each bootstrap iteration *i*, we examined the linear regression of log(*B*) vs log(*P*) and log(*Y*) vs log(*P*), estimating the regression coefficient *b*_*i*_ and its standard error *se*_*i*_. This information was used to draw a random value for the regression coefficient of iteration *i* from a normal distribution with mean *b*_*i*_ and standard deviation *se*_*i*_. The standard deviation of the distribution of these regression coefficients over all bootstrap iterations was then taken as the standard error for the gradient in the field survey. Obtained in this way, the standard error of the gradient captures both the uncertainty of the gradient in the field data, and the uncertainty in the bootstrap.

